# Ecologically and economically sustainable level of timber harvesting in boreal forests – defining the safe operating space for forest use

**DOI:** 10.1101/2024.06.27.600997

**Authors:** Mikko Mönkkönen, Clemens Blattert, Jérémy Cours, Rémi Duflot, Merja Elo, Kyle Eyvindson, Jari Kouki, María Triviño, Daniel Burgas

## Abstract

Planetary-level analyses indicate that we are exceeding the ecological limits. However, we need approaches to implement global sustainability frameworks regionally, using natural resources at levels that allow for their regeneration. We present a framework to define limits beyond which ecosystems are threatened to collapse, to answer how much we can extract from ecosystems, and to manage natural resources for both human and ecosystem wellbeing. We exemplify this approach with the heath forest habitat types in Finland, which provide most of the national timber production. We use the IUCN Red List of Habitats to set favourable reference values for volume of deadwood, proportion of old-growth forest cover and proportion of deciduous trees. Through forest growth simulations and management optimization, we found that the proportion of old-growth forest is the most challenging criteria to be reached only by 2100. This would require not only a larger use of extensive forest management practices than hitherto but also to drastically reduce the maximum economic sustainable harvest level from the current 96% to 60%. By combining threat assessments with ecosystem modelling and management planning, this approach can support regional decision makers to make informed decisions to stay within safe limits of use of natural resources.

## Main

Humans have already exceeded six of the nine Earth System boundaries beyond which humanity can operate safely (i.e., planetary boundaries, *sensu*^1–3^). This overshoot of, for example, climate and biodiversity loss (or biosphere integrity, *sensu*^1^), poses a significant risk for ecosystems and their functions and, ultimately, also for human societies^4,5^. As human appropriation of natural resources and habitat encroachment are the main factors deteriorating the Earth system^6^, we should set boundaries for human activities, yet, defining how this could be done in practice remains a significant challenge. As species always require functional ecosystems and habitats therein, a useful approach to prevent biodiversity loss is to reduce the threat for habitats. IUCN^7^ defines that “an ecosystem is collapsed when it is virtually certain that its defining biotic or abiotic features are lost, and the characteristic native biota is no longer sustained”. To sustain a favourable conservation status of habitats and avoid ecosystem collapse, we must bring human influence into a safe operating space, that is limited by the ecological ceiling and the social foundation—i.e., the unavoidable impacts of human activities to fulfil human needs (following the doughnut economics framework, Fig. 1a;^8^).

**Fig. 1.**
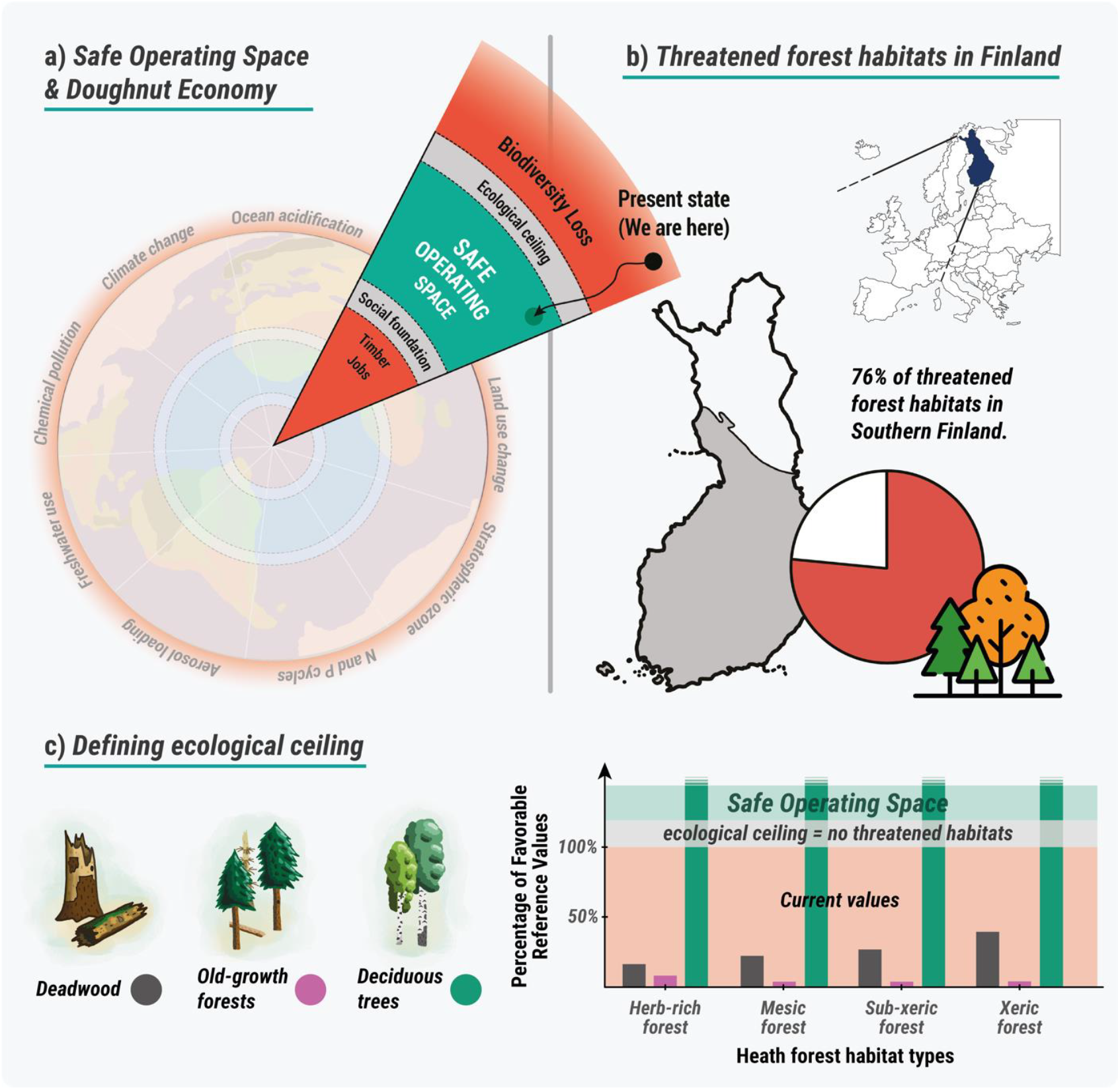
Schematic overview of the framework developed to explore ecologically and economically sustainable timber harvest levels applied to Finland. **a**) Our study combines the frameworks of Planetary Boundaries^1^ and doughnut economics^8^. We evaluate what would be the maximum level of timber extraction, i.e., timber flow and net present value (social/economic foundation) that would still allow critical forest structural features to develop and remain at necessary levels (ecological ceiling). **b**) Case study context of Southern Finland, in grey, covering the hemiboreal, south and middle boreal vegetation zones of Finland. **c**) Current level of the three criteria defining the conservation status of heath forest habitats and, thus, used to define the ecological ceiling (see criteria values in Table 1).

**Table 1.** Reference values for the quality indicators for the four fertility site types in Southern Finland. Values for forests in natural state (no human impact) were calculated multiplying the values for each age class in Kouki *et al*.^53^ by the proportion of land covered by each age class in natural state forests^55^. Values for the 1750 were taken from Kouki *et al*.^53^. Values for the 1960s were taken from the 5^th^ national forest inventory in Finland (NFI5) and current values were obtained from the input data for the simulations (see section Data and forest simulation). The logic to derive favourable reference values (FRVs) is explained in the text.

| Indicator | Fertility site type | Natural state | Ref. 1750 | 1960s (NFI5) | FRV | Current values |
| --- | --- | --- | --- | --- | --- | --- |
| Deadwood volume (m<sup>3</sup>/ha) | Herb-rich | 140 | 56 | 3.3 | 28 | 4.9 |
|  | Mesic | 110 | 38 | 3.7 | 19 | 4.7 |
|  | Sub-xeric | 95 | 29 | 3.7 | 14.5 | 4.4 |
|  | Xeric | 70 | 20 | 3.2 | 10 | 4.3 |
| Proportion of old forest (% cover)* | Herb-rich | 50 | 11 | 0.6 | 5.5 | 0.4 |
|  | Mesic |  | 25 | 4.4 | 12.5 | 0.1 |
|  | Sub-xeric |  | 35 | 1.4 | 17.5 | 0.3 |
|  | Xeric |  | 50 | 0.6 | 25 | 0.5 |
| Proportion of deciduous trees (% volume) | Herb-rich | na | na | 28.5 | 20 | 36 |
|  | Mesic |  |  | 16.9 | 11.8 | 24 |
|  | Sub-xeric |  |  | 8.1 | 5.7 | 12 |
|  | Xeric |  |  | 3.3 | 2.3 | 9 |

\*Threshold ages for old forest: herb-rich 120 years, mesic and sub-xeric 140 years, xeric forests 160 years

Many environmental policy initiatives (e.g., Global Biodiversity Framework) address the challenge by asking “*how much conservation is enough?*” to maintain biodiversity and strive to set the minimum coverage of protected areas (e.g.,^9,10^). In addition, we need to tackle the root causes of biodiversity loss rather than only remedy their symptoms. Biodiversity is lost because we consume natural resources faster than nature can regenerate them^11^. Even when not exceeding the regeneration rate, excessive human appropriation of natural resources is a threat to biodiversity and ecosystem integrity (e.g., functional), as remaining resources (e.g., carbon, nutrients, water, space) may become insufficient. Thus, we should shift the focus and address “*how much resource use is too much?*” and quantify an ecological ceiling that prevents ecosystem collapse. Defining a maximum biodiversity-safe level of natural resource use would provide a clear, practical, and accountable policy target that relates directly to biodiversity loss due to human activities.

Taking the boreal forest ecosystems and forestry sector as an example, we explored how to define an ecological ceiling for the safe operating space. The boreal forest is one of the largest biomes on Earth, contains approximately 45% of the world’s stock of growing timber and accounts for 25-33% of wood-biomass products within the global export market^12,13^. About two-thirds of the biome’s area is actively managed and harvested for timber production, which changes the structure and tree species composition of the forest, and impacts wildlife and ecosystem functioning^14–16^. Particularly in Fennoscandia, protected areas cover only less than 5% of productive forest land and forests are considered primarily a source of biomass^17^. Therefore, habitat and resource availability for species associated with old-growth forests have severely declined. Consequently, forest management is the most prominent threat for biodiversity^18–20^. It is evident that the ecological ceiling of forest ecosystems and their functional integrity *sensu* Rockström *et al*.^21^ have largely been exceeded in Fennoscandia.

Fennoscandian boreal forests are perhaps the most thoroughly studied terrestrial ecosystems of the biosphere. From a global perspective, national economies of the Nordic countries are affluent, stable, and predictable. Therefore, Fennoscandia is an ideal test arena to find out if sustainable forestry, in ecological, economic, and social terms, is an achievable goal. Implementation of the safe operating space consequently implies securing forest habitats while maintaining timber production for the forestry sector. Indeed, the latter represent a significant economic sector, providing raw material, energy, and job opportunities to society (i.e., social foundation, Fig. 1a).

Here, we introduce a new approach to address the question of “*how much is too much?*” and test it by defining an ecological ceiling for human forest use, that is, a maximum timber harvest level below which forest ecosystems are no longer threatened. We apply this approach to the hemiboreal, south and middle boreal heath forests in continental Finland (hereafter Southern Finland, *sensu* ^22^) using the recent Red-List assessment of forest habitat types (Fig. 1b;^22^). Specifically, we ask (i) how fast the forest habitat types can be removed from the IUCN Red List, (ii) how forests should be managed to achieve this goal, and finally, (iii) what is the maximum annual harvest to stay in the safe space. To answer these questions, we use the IUCN Red-List criteria^22^, which define criteria for endangerment of forest habitat types, and identify the level of those criteria that should be reached to remove the forest habitats from the risk of collapse, i.e., Favourable Reference Value (FRV, Fig. 1c). Then, we use forest growth simulations and a multi-objective optimization framework^23^ to define the time, the optimal management combination, and the maximum harvesting level needed to reach the FRVs (Fig. 2).

**Fig. 2.**
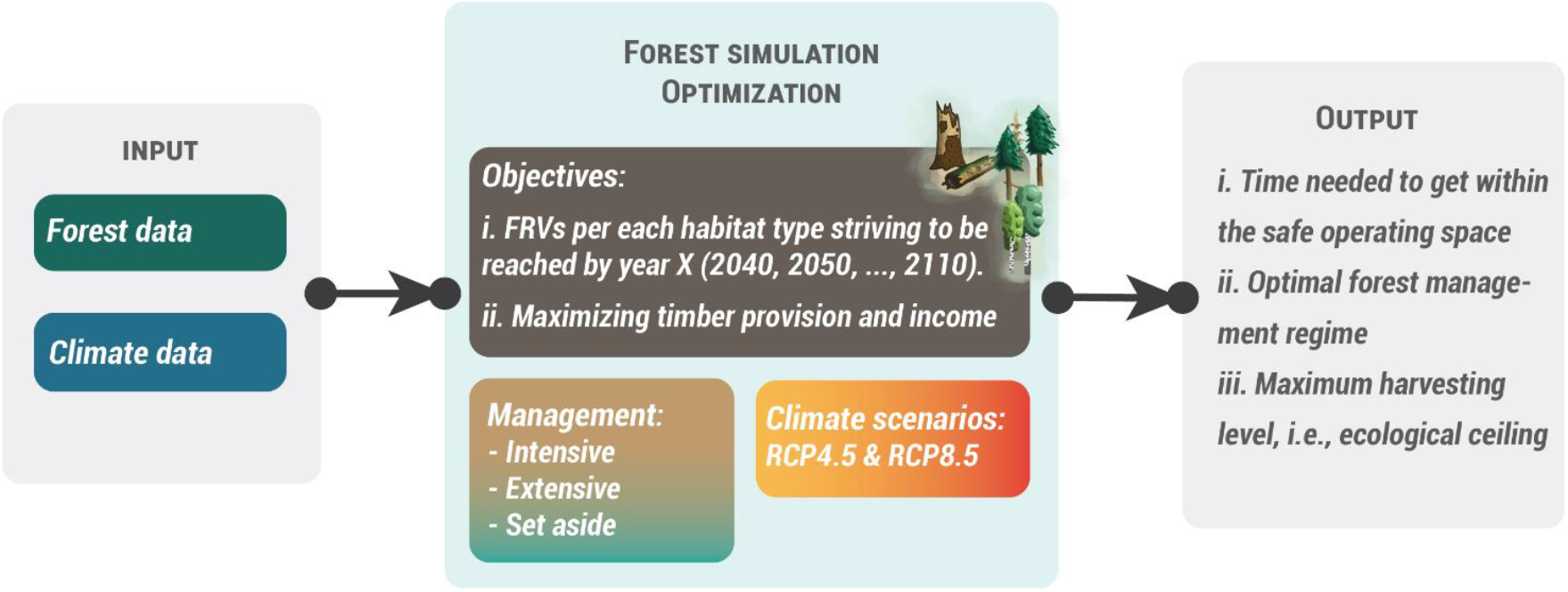
Workflow used for the analyses and main answers we expect with the outputs of this framework. FRVs stands for Favourable Reference Values.

## Results

### How much time will it take to reach favourable reference values for all forest habitat types?

The time needed to render the habitat types not threatened ranges from 40 to 80 years (2060-2100), depending on the habitat type (Fig. 3). In xeric forests (representing 3% of forest area in Southern Finland^24^), it may take till 2100 before all criteria can reach their favourable reference values (FRVs). Herb-rich forests (23% of forest area) can reach FRV already by 2060, mesic by 2070 (50 % of forest area) and sub-xeric forests 20 % of forest area) by 2080. The main limiting criteria affecting the time needed to achieve a favourable status was the proportion of old-growth forest area, except for herb rich forest for which the limiting factor was the volume of deadwood (Fig. 3, see Table 1 in the Methods section). The required proportion of deciduous trees from total timber volume is swiftly achievable across the forest types. To project the development of forests, we used the intermediate climate change Representation Concentration Pathway (RCP) 4.5, which approximates our current climate pathway^25^. Sensitivity analyses with the “business as usual” high-emission scenario RCP8.5 did not show significant changes in the timing or criteria limiting the achievement of favourable status (Supplementary Material Fig. S1).

**Fig. 3.**
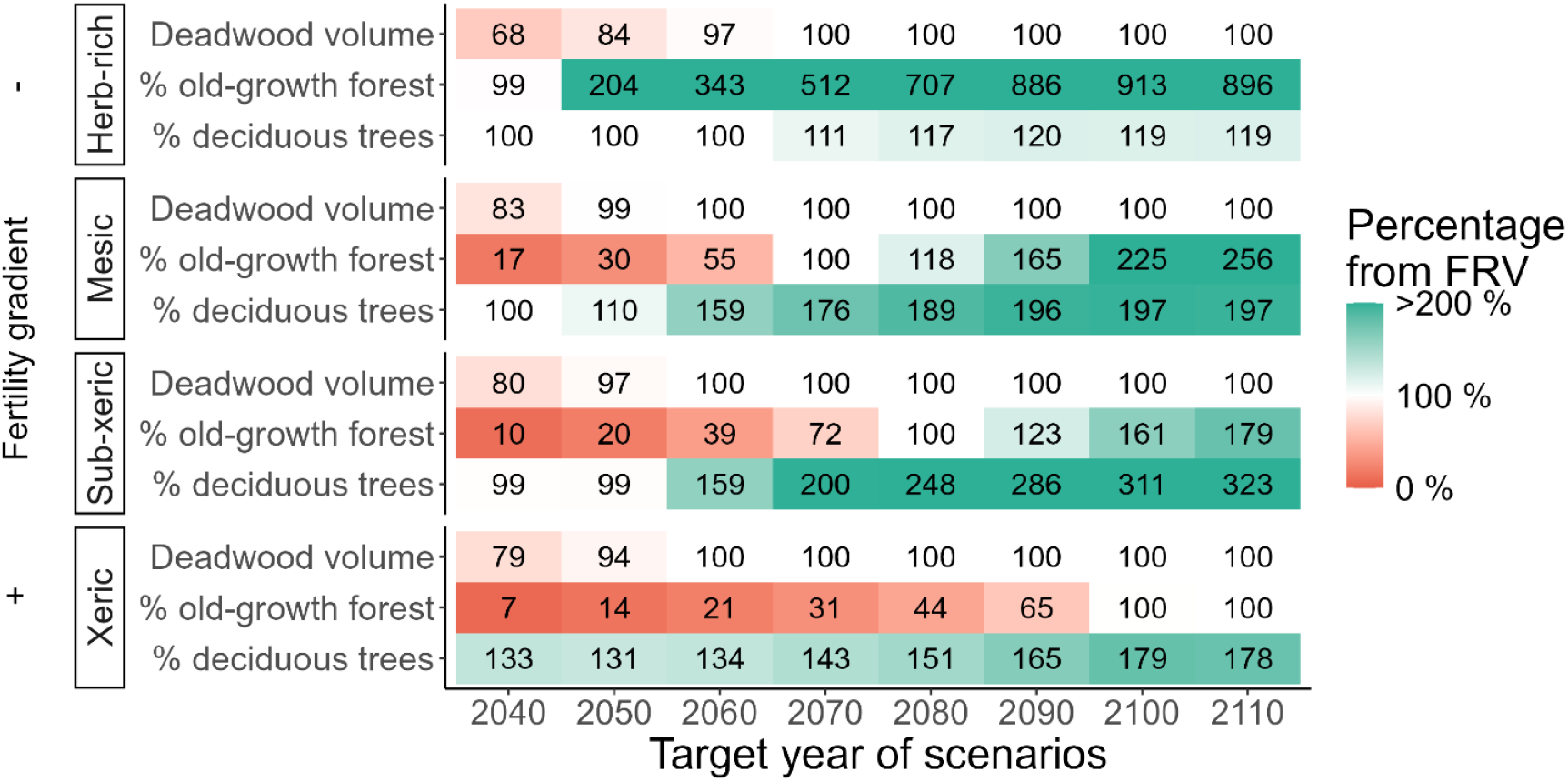
Possibility to achieve and exceed the favourable reference values (FRV) in Southern Finland for the four main habitat types of heath forest (herb-rich, mesic, sub-xeric and xeric). The horizontal axis represents the target year when the FRVs were strived to be reached. Values for alternative criteria are expressed as percentages from the FRVs (see Table 1), with red colour indicating that the target has not yet been achieved, white that the target has been achieved and green that the target has been surpassed.

### How forests should be managed and what is the maximum harvest level to stay within a safe operating space?

The faster we strive to render forest habitat types not threatened, the more we need to rely on setting aside forests from timber harvesting (Fig. 4a). Targeting 2100, the earliest year when all forest habitat types can reach favourable reference values, we would need to set aside (i.e., no management) more than one third of forests. This is challenging as currently less than 6% of forest land in Southern Finland is protected^26^. The optimal combination of management categories also included 40% of intensive management where the focus is more on timber production, and 25% of extensive management also targeting biodiversity conservation and other ecosystem services than timber (Fig. 4a, and Supplementary Material Fig. S2 for management sub-categories). There was variation among the forest habitat types in the optimal combination of management categories. In general, on the more productive habitat types (herb-rich and mesic forests) set aside was a more prominent option, while on less productive habitat types (sub-xeric & xeric forests) extensive management gained more prevalence (see Supplementary Material Fig. S3 for the proportion of management showed by habitat type categories).

**Fig. 4.**
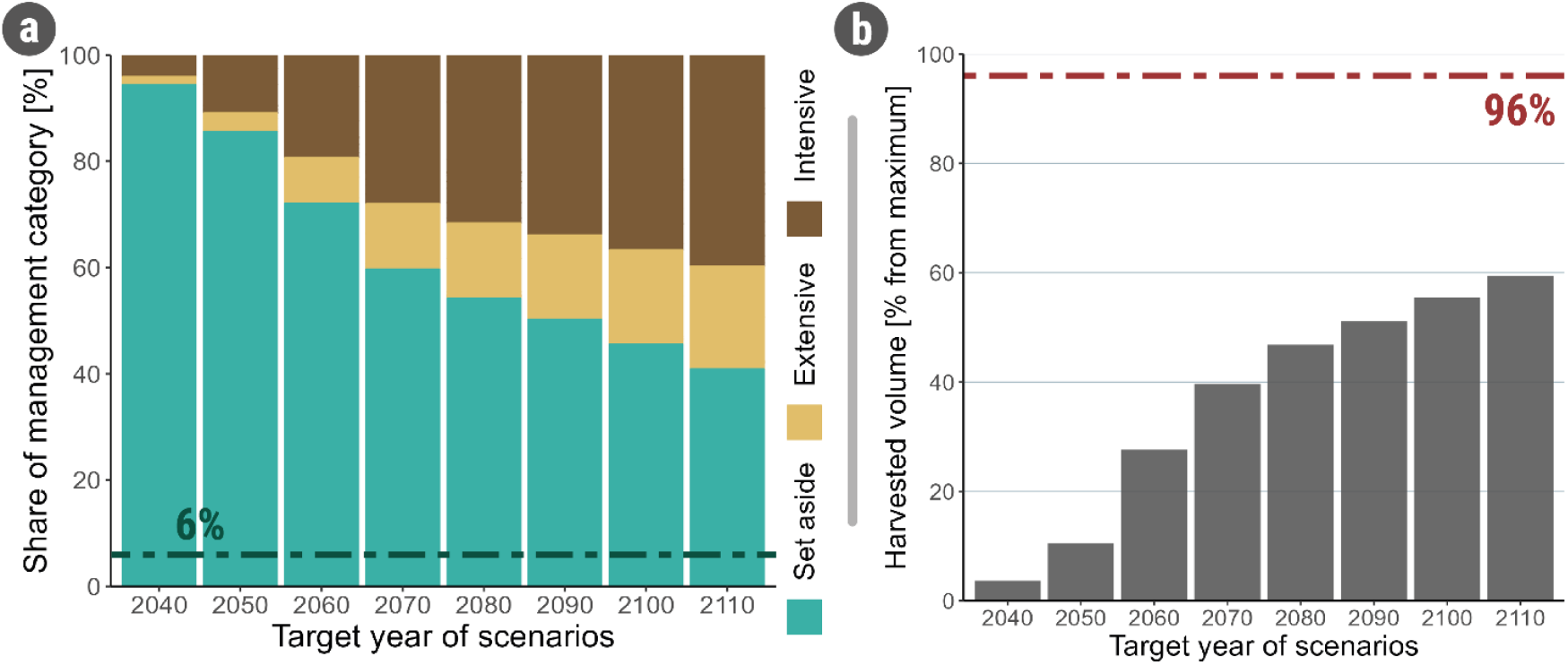
(a) Proportion of management categories and (b) maximum annual timber harvest level that would maximise the favourable reference values in all four forest habitat types. The horizontal axis represents the target-year when the FRVs were strived to be reached (but might not be reached, see Fig. 3). In (b), the vertical axis shows the harvest level relative to the maximum economically sustainable annual harvesting potential (i.e., without ecological constraints). The dashed horizontal line represents the current level of set-aside forest land (a), and the average harvest level between the years 2016-2021 (b) in Southern Finland^27^.

The sooner we want to reach FRVs, the more we should give up on the maximum economically sustainable annual harvesting potential, i.e., the maximum annual harvest allowing for a constant timber production over time (also known as even flow of timber) (Fig. 4b). If we set 2100 as the target year, when all forest habitat types can reach a favourable status, the ecological ceiling is a harvest level of 58% of the maximum economically sustainable annual harvesting potential. This is considerably less than the realized recent harvest level in Southern Finland, that is 96% (Fig. 4b, ^27^).

## Discussion

Our economies are not external to nature but embedded within it^11^. Building on this premise, we operationalized the planetary boundaries and doughnut economy frameworks at large spatial scale and defined a maximum biodiversity-safe level of harvesting timber from a European boreal region. Maximum harvest level was estimated so that it does not exceed the capacity of the forest ecosystems to supply biodiversity-critical habitat characteristics. Results indicate that harvesting should be capped well below current levels, but the exact level depends on how fast we want to enter the safe operating space. Setting a realistic target year of 2100 requires utilizing at most 60% of the annual harvesting potential. Curbing forest harvesting to this level creates a challenge for the Fennoscandian forest industry, which currently is largely based on producing large quantities of pulp and sawn timber for exporting products. The challenge can be tackled by developing regenerative business models^28^, which emphasize higher value-added wood products that provide livelihoods for the entire value-chain relying on wood biomass. More profoundly, resolving this challenge will require moving away from a life frame where nature is considered a mere provider of resources, to partnerships with nature that focus on nature’s life-supporting processes^29^.

Simultaneously with the ecological ceiling, we evaluated the social foundation of the Finnish forest use as the maximum annual timber flow and monetary benefits (net present value) that are possible to achieve under the ecological ceiling. This follows the current societal values, which emphasize instrumental values and where forest resources are seen as a provider of job opportunities, economic value, and raw materials to fulfil human needs^30^. In affluent postindustrial societies, such as in Fennoscandia, we see an increasing endorsement of relational and intrinsic values such as seeing nature as a part of one’s meaningful life and much more than means to an end. Such values are typically tied to local context, and thus, implementing the idea of staying within the safe operating space requires local deliberation and planning—e.g., on how to sustain livelihoods with less resource extraction in a just manner.

Staying within a safe operating space requires a constant flow of forest biomass, which, according to our results, is best achieved with a diverse combination of management regimes to ensure a diversity of habitats^31,32^ and the flow of timber and non-timber ecosystem services from forests^33^. Our results suggest that high shares of both protected areas supporting biodiversity conservation, and intensively managed areas—maximizing timber production and economic benefits—are needed, complemented by extensive management. The utility of such functional zoning outcome is evident^34,35^. This requires a big change in current management practices, where intensive clear-cut forestry heavily dominates in Fennoscandia, by increasing adoption of alternatives such as continuous cover forestry (see Supplementary Material Fig. S2).

Among the biodiversity indicators, achieving the favourable reference value for old-growth forests is the largest challenge. More than a century of intensive forest management has shifted the forest composition towards young forests^36^ and recovering a high proportion of old-growth forest will require a long time. This result highlights the need to swiftly start actions for biodiversity as recovery processes and formation time of most critical microhabitats in forest ecosystems are slow. Only by acting now we can reduce any further, potentially irreversible biodiversity losses^37^. The primary focus should be on safeguarding the remaining old-growth forests, in accordance with the EU biodiversity strategy for 2030^9^. However, old-growth forests are not only characterized by tree age but also by their structural features that are important for biodiversity and human well-being^38,39^. Hence, there exists the possibility of actively restoring deadwood and tree-related microhabitats to speed up the recovery process^40^.

Deadwood is a critical resource for species in boreal forests^41^ and a good indicator of forest biodiversity^42^. In boreal Fennoscandia, 20–25% of the forest-dwelling species are dependent on deadwood habitat and they constitute 60% of red-listed species^43^. The FRVs for deadwood volume can be reached relatively quickly in all forest types, suggesting the potential for relatively rapid recovery of deadwood-dependent biodiversity. However, production of large deadwood pieces, sheltering peculiar communities^44^, should need extra-time. This effectively requires more set asides^36^, which would eventually result in more old-growth forests, as well.

Further enhancements of this approach could entail adding ecological constraints into simulations as well as considering social factors. For example, our results remained qualitatively similar under different climate change scenarios, but this could change by including natural disturbances, which have increased and will continue to increase in frequency, intensity and spatial extent due to climate change^45,46^. Disturbances may have contrasting effects on FRVs: they contribute in creating deadwood and certain tree-related microhabitats if systematic salvage logging of damaged trees is avoided^36,47,48^, while severe disturbances can also reduce the area of old-growth forests^49,50^. Ultimately, cumulative and interactive effects of human and atypical natural disturbances may lead to an ecosystem shift into a compromised functional state^45,51^. Because reaching a favourable conservation status for heath forest habitats in Finland will take several decades, the chance of facing extreme disturbance events is very high, further delaying the achievement of targeted proportion of old-growth forest. Hence, the more ambitious we are initially, the more likely we are to halt the biodiversity crisis.

In our study we consider the ecological and economic limits of the forest ecosystems to identify a safe operating space. Future studies should go one step further and quantify both safe and just Earth system boundaries^21^. Thus, a just transformation of the forest sector requires acknowledging the multiple needs and views of forest stakeholders and understanding the dynamics of the entire forest social-ecological system transition.

## Methods

We used a large forest dataset representing the hemiboreal, south and middle boreal vegetation zones in Finland (Fig. 1). We focused on the four main fertility site types of heath forest: herb-rich, mesic, sub-xeric, and xeric heath forest. These four site types represent together over 96% of the productive forest of the focal regions according to the latest National Forest Inventory (NFI12). Forested peatlands were excluded because a large proportion of them have been drained for forestry purposes, which is the main criteria defining their conservation status, i.e., not directly related to harvest level. We also excluded the northern boreal vegetation zone, which contains 28% of heath forests in Finland, because a large proportion of heath forests there are either barren heaths (very low timber production) and/or currently protected. Less than 5% of harvested timber volume in Finland originates from forests in the North boreal zone, and thus, excluding these forests has a small effect on required harvesting levels at the national scale. Moreover, most threatened forest-dwelling species in Finland have Southern distributions^20,37^.

### Favourable reference values for defining an ecological ceiling

The recent assessment of threatened habitat types in Finland^52^ followed IUCN criteria, which consider changes in the quantity of the habitat type, changes in abiotic and biotic quality, and rarity. The assessment shows that three-quarters of the Finnish forest habitat types are threatened^22^; herb-rich and mesic forest habitats are classified as vulnerable (VU), i.e., at a high risk of collapse, and sub-xeric and xeric forest habitats are endangered (EN), i.e., at a very high risk of collapse.

Most of the heath forest habitat types were assessed as threatened according to changes in their abiotic and biotic quality (IUCN criteria D). However, the decline in area (IUCN criteria A) was the most important reason for old heath forests to be threatened^22^. We adopted three quality indicators, which translate the IUCN criteria into measurable characteristics, from Kouki *et al*.^53^: deadwood volume, the proportion of old forest cover, and the proportion of deciduous trees relative to the total tree volume. These indicators are considered critical features also for red-listed species in Finnish boreal forests^20^. Following the logic of the greatest risk of loss and following the IUCN guidelines, Kouki *et al*.^53^ used the condition of forests in 1750 as a reference point, except for the proportion of deciduous trees for which the situation in the 1960s was the reference (the value in 1750 is not available). The 1750 time point represents the earliest onset of industrial-scale exploitation of ecosystems^54^ and is justified in the IUCN Red List of Ecosystems^7^. According to the IUCN criteria D, a habitat type (fertility site type in our study) is considered threatened if the indicator value has decreased more than 50% (30% for the proportion of deciduous trees) from its reference value in 1750 (1960s for the proportion of deciduous trees). We followed this logic to address what is the required improvement in the quality indicator values that will render the heath forest habitat types not being at risk of collapse (i.e., not threatened). That is, for a currently threatened habitat type to become not threatened, the value of an indicator must improve to a level that is at least 50% of the reference value in 1750 (area of old forest and volume of deadwood), or 70% of the reference value in the 1960s (proportion of deciduous trees) and remain at that level or above ever since. We consider this level of an indicator as a favourable reference value (FRV, Table 1).

We used IUCN criteria to estimate favourable reference values. Another option would have been to derive the FRVs from literature on species distributions and population persistence as functions of critical structural features. Although there is much variation in species response to variation in critical features^56^, literature suggests that when resource availability decreases by 80-90% from the natural state, native biota become depauperated due to local extinctions^57–59^. This approach would result in similar FRVs to what we had (Table 1). Our FRVs for deadwood volume also largely correspond to critical ecological threshold values for deadwood-dependent organisms in boreal forests^60,61^.

### Data and forest simulations

To obtain a representative sample of the initial forest state for forest growth simulations, we used two data sources. We obtained private forest owners’ detailed stand-level information from 2016 from the public database www.metsaan.fi of the Finnish Forest Centre. This data was complemented by the Multi-source National Forest Inventory from 2015, to cover areas not privately owned^62^. Both data sets were sampled along the regional and temporal systematic clusters following the design of the 11th Finnish National Forest Inventory (see ^23^). We selected only data points within heath forest of the hemiboreal, south and middle boreal vegetation zones in continental Finland.

We performed the forest simulations using the open-source forest simulator SIMO^63^. SIMO simulates individual tree growth, mortality, and regeneration for even-aged^64^ and uneven-aged (i.e., continuous cover forestry) boreal forests^65^. We simulated forest dynamics and management under two climate trajectories; representative concentration pathways RCP4.5 and RCP8.5. Climate variables driving stand growth and soil dynamics (mean and amplitude of temperature, CO_2_ concentration, precipitation) were based on Lehtonen *et al*.^66^, and the climate data of the Canadian Earth system model CanESM^67^. The impacts of climate on tree growth are introduced into the calculation of volume growth, and further allocated between diameter and height growth based on the models of Matala *et al*.^68^.

For each forest plot, we simulated 29 management regimes in five-year periods over 100 years, with the exact number of management alternatives applied depending on the initial conditions of each individual stand (i.e., dominant stand height, basal area, site type, and stand age). The regimes provided a diverse set of management alternatives for each stand, from which the optimization afterward selected an ideal management based on the management effects on optimization objectives (see optimization section below). We grouped the regimes into three classes based on the harvest level: Intensive, Extensive, and Set aside (Table 2).

**Table 2.** Summary table of the simulated management regimes for Finland and their allocation to the six management classes. The classification to intensive and extensive regimes follows Blattert *et al*.^35^. BAU stands for business-as-usual rotation forestry; CCF for continuous cover forestry.

| Management class | Abbreviation | Description |
| --- | --- | --- |
| Intensive managements | *BAU* | Rotation forestry, no thinnings prior clearfelling |
|  | *BAU F* | = *BAU*, with fertilization |
|  | *BAU -5* | = *BAU*, with 5 years shorter rotation age |
|  | *BAU -5 F* | = *BAU -5*, with fertilization |
|  | *BAU + 5* | = *BAU*, with 5 year extended rotation age |
|  | *BAU w thin* | Rotation forestry, with thinnings prior clearfelling |
|  | *BAU w thin B* | = *BAU w thin*, with increased broadleaf planting |
|  | *BAU w thin F* | = *BAU w thin*, with fertilization |
|  | *BAU w thin -5* | = *BAU w thin*, with 5 years shorter rotation age |
|  | *BAU w thin -5 F* | = *BAU w thin -5*, with fertilization |
|  | *BAU w thin +5* | = *BAU w thin*, with 5-year extended rotation age |
|  | *BAU w thin +5 B* | = *BAU w thin +5*, with increased broadleaf planting |
|  | *BAU w thin GTR* | = *BAU w thin*, with 30 retention trees or 30m<sup>2</sup> per ha left |
|  | *BAU w thin GTR B* | = *BAU w thin GTR*, with increased broadleaf planting |
|  | *BAU w GTR* | = *BAU*, with 30 retention trees or 30m<sup>2</sup> per ha left |
|  | *CCF 1* | Thinning from above, basal area threshold -3 m<sup>2</sup>/ha |
|  | *CCF 2* | Thinning from above, basal area threshold +/-0 m<sup>2</sup>/ha |
| Extensive managements | *CCF 3* | Thinning from above, basal area threshold + 3 m<sup>2</sup>/ha |
|  | *CCF 4* | Thinning from above, basal area threshold + 6 m<sup>2</sup>/ha |
|  | *BAU w/o thin* | Rotation forestry, no thinnings prior to or after clearfelling |
|  | *BAU w/o thin -20* | = *BAU w/o thin*, with 20 years shorter rotation age |
|  | *BAU w/o thin -20 F* | = *BAU w/o thin -20*, with fertilization |
|  | *BAU + 15* | = *BAU*, with 15 years extended rotation age |
|  | *BAU w thin +15* | = *BAU w thin*, with 15 years extended rotation age |
|  | *BAU w thin +15 B* | = *BAU w thin +15*, with increased broadleaf planting |
|  | *BAU +30* | = *BAU*, with 30 years extended rotation age |
|  | *BAU w thin +30* | = *BAU w thin*, with 30 years extended rotation age |
|  | *BAU w thin +30 B* | = *BAU w thin +30*, with increased broadleaf planting |
| Set aside (SA) | *SA* | No management, only growth and mortality |

### Optimization

To quantify the potential ability of the forest to reach the specific FRVs at a specific time period, we applied a systematic optimization framework. This framework is based on the MultiOptForest framework^69^ which relies on the application of an achievement scalarizing function ^70^ and the epsilon constraint method^71^. This allowed evaluating the possibility to reach the specific targets, and the economic timber availability possible once the maximum potential (or targeted) habitat indicators are reached. The general optimization problem can be formulated as:

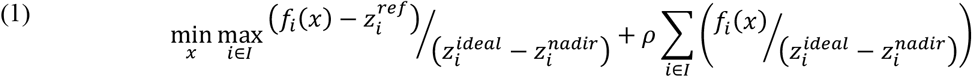

subject to:

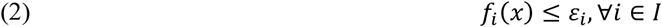

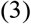

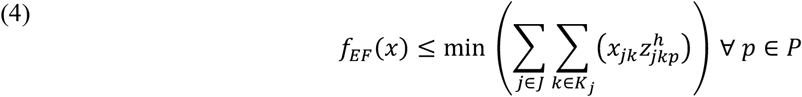

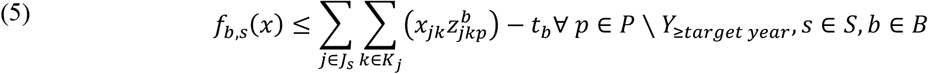

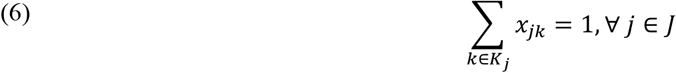

where *I* is the set of objectives; *f*_*i*_(*x*) is the obtained value for the specific objective; 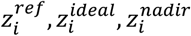 are the target, ideal and nadir values for the specific objective; *ρ* is an arbitrary small constant; *ε*_*i*_ is the epsilon constraint target for the specific objective; *P* is the set of periods under consideration; *J* is the set of forest plots; *K*_*j*_ is the management options available for stand *j*; 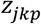 is the value provided for each stand, management action at the specific period, with superscripts *i, e, h and b* respectively referring to net income, end stand value, timber harvest quantity and, FRVs; *x*_*jk*_ is the decision to conduct management action *k* to stand *j*.

The optimization problem starts with equation (1), where the objective function is a multi-objective optimization, formulated as an achievement scalarizing function. The epsilon constraint is formulated in equation (2), which requires the objective to be greater than the specified value (*ε*_*i*_). Equation (3) evaluates the net present value of the problem, with equation (4) calculates the minimum timber harvest value across all periods. Equation (5) calculates the FRVs, striving to meet the target value at a specific period, and exceeding it for all other periods.

To assess when the indicators are possible to be attained, we performed a re-iteration of the optimization striving to achieve the FRVs targets at the beginning of each decade. We first ran the optimization without any epsilon constraints (*ε*_*i*_) and set the targets 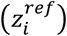 for the achievement scalarizing functions to the specific FRVs desired. We then ran the optimization by setting the obtained values for the FRVs as epsilon constraints (requiring the specific indicator value at the specific year), and re-ran the optimization three times: once were we strived to maximize NPV, to maximize the minimum periodic timber harvested across all time periods exceeding the targeted year, to maximize both the NPV and minimum periodic timber harvested across all time periods exceeding the targeted year.

## Acknowledgements

This work was partly funded by the Strategic Research Council at the Academy of Finland (Mönkkönen 358473). J.C., M.T. and R.D. were supported by the Kone Foundation (application 202206136 and 202105759).We would also like to thank Kaisa Junninen for the help with threat assessments of forest habitats.

## Author contributions

M.M., C.B., R.D., M.E., K.E., M.T. and D.B. conceived and designed the study, performed the experiments, and analysed the data. C.B., K.E. and J.K. contributed materials/analysis tools. J.C. contributed with visualizations and analyses. All authors contributed to the writing of the manuscript.

