## Supplementary material for "Ecologically and economically sustainable level of timber harvesting in boreal forests – defining the safe operating space for forest use"

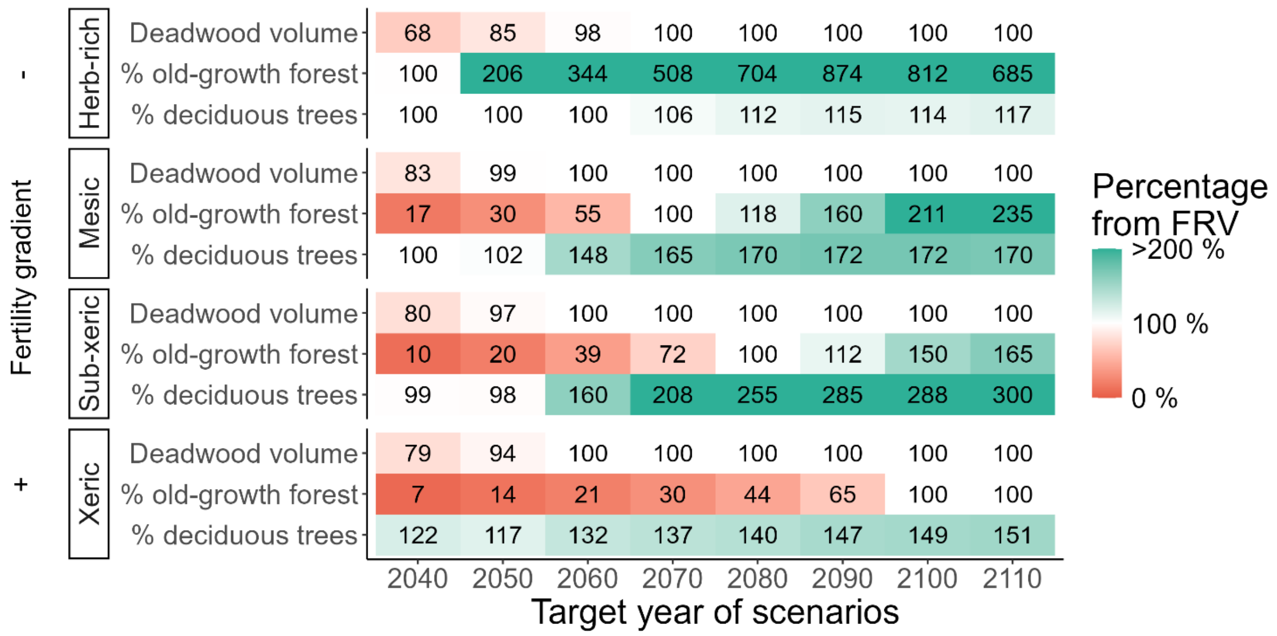


**Fig. S1 | Possibility to achieve the favourable reference values (FRVs) in Southern Finland for the four main fertility classes in heath forest (herb-rich, mesic, sub-xeric and xeric) under climate scenario RCP8.5**. The horizontal axis separates cost-effective management plans aiming to achieve the FRVs by each specified target year. Values are expressed as percentages from the target FRVs, with red colour showing that the target has not yet been achieved, white that the target has been achieved and green showing that the target has been surpassed.

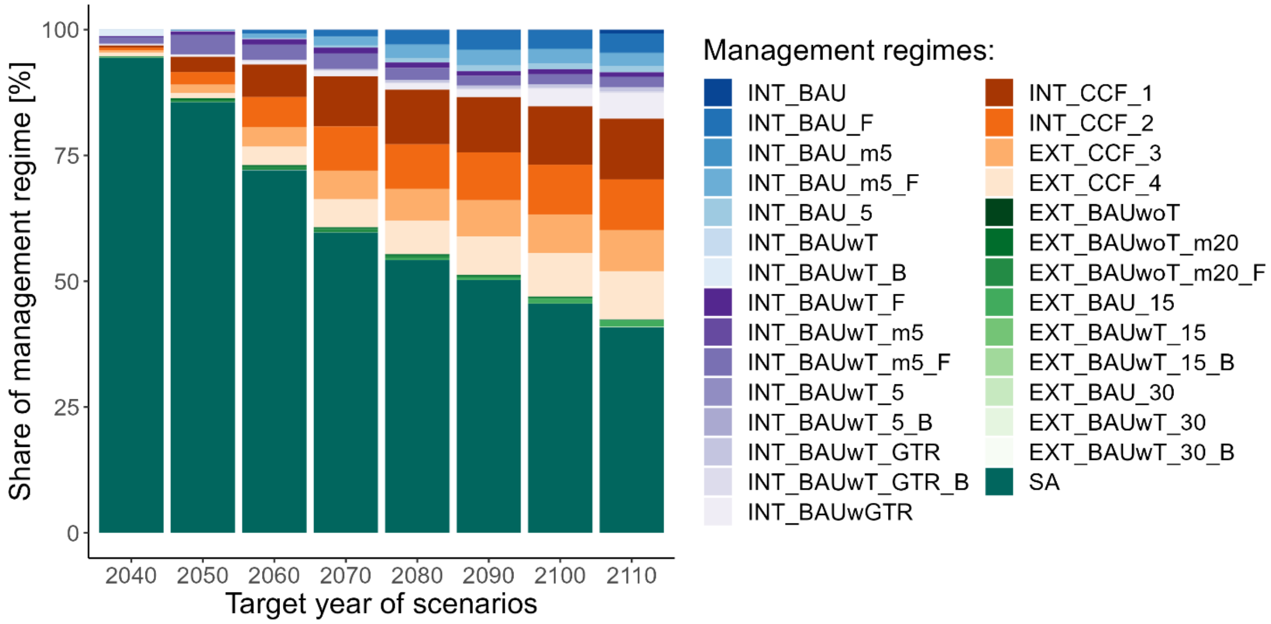


**Fig. S2 | Change in detailed management regimes for each target-year scenario.** Each bar represents a scenario where we tried to achieve the FRVs by a specific year and then stay above the FRVs while maximizing the net present value and timber harvesting volumes. Descriptions of management abbreviations can be found in Table 2 in the methods section.

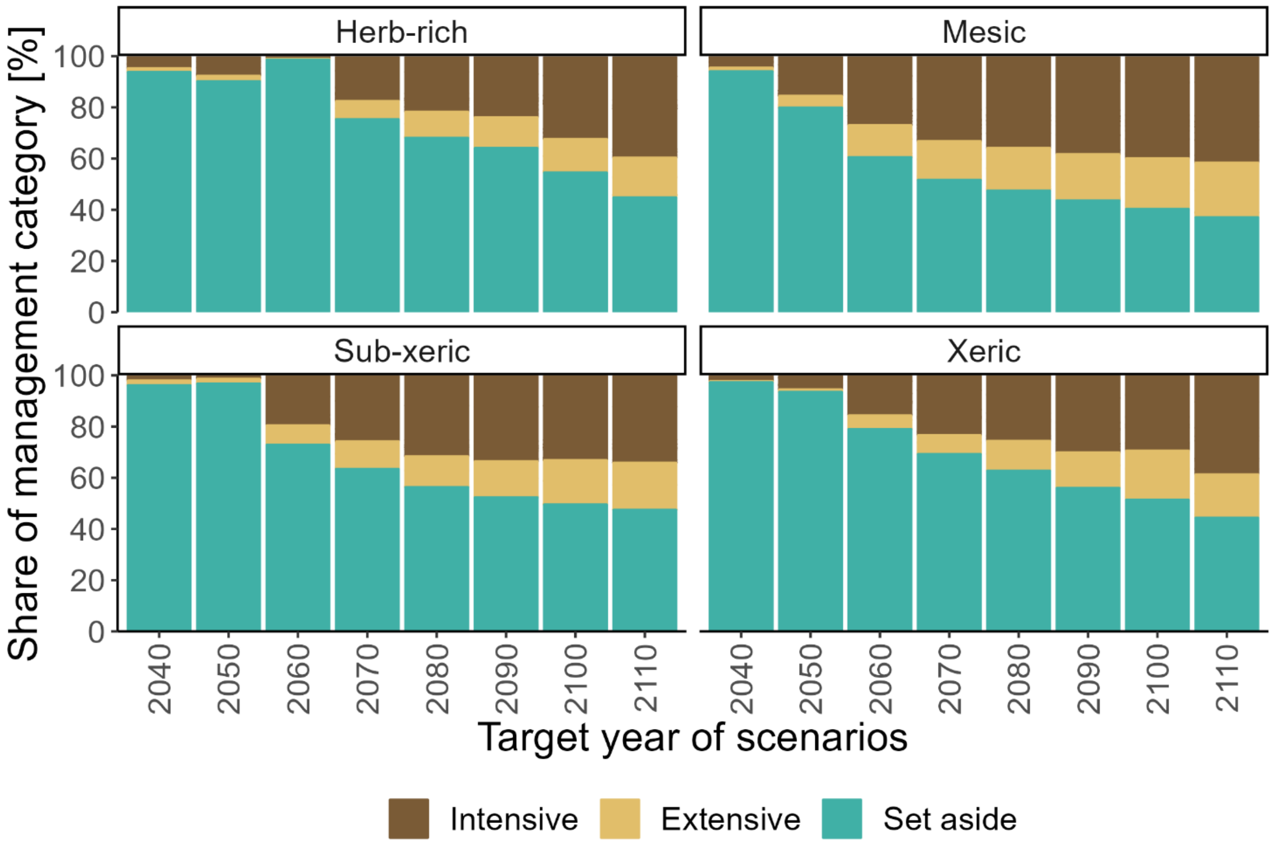


**Fig. S3 | Change in management categories at each habitat type for each target-year scenario.** Each bar represents a scenario where we tried to achieve the FRVs by a specific year and then stay above the FRVs while maximizing the NPV and timber harvesting volumes.
